# A Recombinant Antibody For Tracking Murine Gammaherpesvirus 68 Uracil DNA Glycosylase Expression

**DOI:** 10.1101/2023.05.17.541089

**Authors:** Yunxiang Mu, Joshua B. Plummer, Monika A. Zelazowska, Somnath Paul, Qiwen Dong, Zaowen Chen, Laurie T. Krug, Kevin M. McBride

**Author notes:** Duchossois Family Institute, University of Chicago, Chicago, IL 60637, USA.

## Abstract

Antibodies are powerful tools to detect expressed proteins. However off-target recognition can confound their use. Therefore, careful characterization is needed to validate specificity in distinct applications. Here we report the sequence and characterization of a mouse recombinant antibody that specifically detects ORF46 of murine gammaherpesvirus 68 (MHV68). This ORF encodes the viral uracil DNA glycosylase (vUNG). The antibody does not recognize murine uracil DNA glycosylase and is useful in detecting vUNG expressed in virally infected cells. It can detect expressed vUNG in cells via immunostaining and microscopy or flow cytometry analysis. The antibody can detect vUNG from lysates of expressing cells via immunoblot under native conditions but not denaturing conditions. This suggests it recognizes a confirmational based epitope. Altogether this manuscript describes the utility of the anti-vUNG antibody and suitability for use in studies of MHV68 infected cells.

## Introduction

All Herpesviruses encode an enzymatically active homolog of uracil DNA glycosylase (UNG) (Geoui et al., 2007; Krusong et al., 2006; Lu et al., 2007; Minkah et al., 2015; Ranneberg-Nilsen et al., 2008). Mammalian UNG is the major enzyme involved in removing genomic uracils through the base excision repair (BER) pathway. Uracil in DNA arises from spontaneous or enzymatic cytosine deamination and misincorporation during replication. The U:G mismatches that result from deamination are 100% mutagenic and will result in C to T transition mutations if replicated over. However, uracil is efficiently removed by UNG, which triggers high-fidelity BER, resulting in essentially error-free repair and genomic integrity (Krokan et al., 2014). The specific contributions of herpesvirus and mammalian UNGs to the viral lifecycle and pathogenesis is not completely understood.

Murine gammaherpesvirus 68 (MHV68) is a natural virus that has been utilized in mouse infection models. MHV68 is related to the human gammaherpesvirus Epstein-Barr virus and Kaposi Sarcoma herpesvirus. A transgenic version of MHV68 expressing histone H2B fused to EYFP (MHV68-YFP) allows the detect of infected cells in culture or live mice and serves as a tractable infection model (Collins and Speck, 2012). In the MHV68 virus, open reading frame 46 (ORF46) encodes the viral homolog of UNG (vUNG). MHV68 with a stop codon in the ORF46 (vUNG) is defective in replication and reactivation (Minkah et al., 2015). While MHV68 with ORF46 catalytic domain mutations that cause enzymatic deficiency, does not display replication defects (Dong et al., 2018). Only with a combination of catalytically defective ORF46 and ORF54 encoding the viral dUTPase was a replication defect observed. This suggests a role for the MHV68 vUNG beyond its catalytic function. To study the role of the protein in viral replication and pathogenesis we created an antibody against MHV68 ORF46/vUNG. Here we provide data for the characterization of the antibody and demonstrate its utility to detect vUNG in infected cells.

## Materials and Methods

### Production of Recombinant Proteins

Recombinant murine UNG or vUNG encoded by MHV68 ORF46, was expressed with an N-terminal polyhistidine tag from pET-duet in *E. coli* BL-21 DE3. 1L of *E. coli* production media (Super Broth + 100ug/ml Ampicillin + 100uM FeCl3 + 0.4% glycerol). Cultures were inoculated in a baffled shaker flask with a starter culture and incubated at 37degC with 300rpm agitation. The OD_600_ was read every 30 minutes; once the culture began to approach stationary growth phase the culture temperature was reduced to 18°C and induced with 0.1mM IPTG overnight. Cultures were harvested, lysed by freezing and sonication, and centrifuged at 10kxg for 20 minutes at 4°C. Clarified supernatants were purified using HisTrap excel resin on ÄKTA FPLC or using drip columns of TALON resin. Eluted fractions were separated by SDS-PAGE and stained with coomassie blue to determine purity and amount.

### Antibody production and purification of Recombinant Monoclonal Antibodies

The Recombinant Antibody Production Core (RAPC) at University of Texas MD Anderson Cancer Center was used to create the recombinant antibody characterized in this study. In brief the RAPC used a pipeline to isolate antigen specific plasma cells from immunized C57Bl6/J mice, flow sort single antigen specific cells and clone matching immunoglobulin heavy and light chain variable regions into plasmids that encoded mouse IgG1 constant region. For recombinant antibody production, ExpiCHO Expression System was used (Gibco). Cells were cultured according to manufacturer’s recommendations in ExpiCHO Expression Medium. After 5 days the media was collected 0.45um filtered and purified using HiTrap protein G resin on ÄKTA FPLC (GE). Eluted fractions were quantified using Nanodrop 2000 (Thermo). Polyclonal sera reactive against vUNG (anti-ORF46) was previously described (Dong et al., 2018).

### Characterization of Recombinant Monoclonal Antibodies

Purified antibody was tested by direct ELISA. For recombinant protein antigens ELISA plates (Nunc MaxiSorp) were coated with 10ug/ml recombinant antigen in PBS overnight at 4degC, washed with TBS + 0.1% tween 20 (TBS-T), and blocked with 1% BSA in TBS-T. In both cases, dilutions of purified recombinant antibody as well as positive control polyclonal serum were incubated overnight at 4°C, washed with TBS-T, probed with anti-mouse IgG-AP, washed with TBS-T, and developed with pNPP substrate (Sigma). 405nm absorbance was determined at 10, 30, and 60 minutes using a FLUOStar Omega Plate Reader (BMG Labtech).

For immunofluorescent staining of fixed cells, transiently transfected 293 cells were grown on either gelatin-coated glass coverslips for microscopy or 10cm culture dishes for flow cytometry. Cells to be analyzed by flow cytometry were harvested into 15ml conical tubes by trypsinization prior to staining. Cells were subsequently fixed in 10% buffered formalin and permeabilized with 0.1% triton X100 in TBS. Primary antibodies were used at 5ug/ml in 2% normal goat serum in TBS-T (blocker) to stain target antigen overnight at 4°C. Primary antibody was detected with anti-mouse H+L-AlexaFluor 647 secondary antibody at 1ug/ml in blocker overnight at 4 C. Cells to be analyzed by microscopy were counterstained with DAPI. Samples were imaged with either a Zeiss 510 meta confocal laser scanning microscope or BD LSRFortessa flow cytometer. MHV68-H2bYFP (Collins and Speck, 2012) and MHV68-H2bYFP-ORF46-stop (ΔUNG) (Minkah et al., 2015) as well as infection of cells have been described (Dong et al., 2018).

Purified antibodies were also tested by western blot. 100ng of recombinant antigen and 50ug cell lysate was separated on 12% SDS-PAGE, transferred to PVDF, and blocked with 3% BSA in TBS-T. The blot was incubated with 1ug/ml purified antibody for 2 hours at room temperature with 3% BSA in TBS-T, washed with TBS-T, and probed with anti-mouse IgG-HRP for 1 hour at room temp in 3% BSA in TBS-T. The blot was washed again in TBS-T, developed with West Pico ECL (Thermo), and scanned on an ImageQuant LAS 500 (GE).

## Results and Discussion

A recombinant mouse antibody against the viral uracil DNA glycosylase (vUNG) of murine gammaherpesvirus 68 (MHV68) was produced by the recombinant antibody production core (RAPC) at the University of Texas MD Anderson Cancer Center. The sequence of the antibody heavy and light chain variable domain sequence is displayed in Figure 1. The antibody was recombinantly expressed as a murine IgG1 isotype in CHO cells and purified with protein A purification. The viral UNG shares high sequence identity with mammalian UNGs, particularly in the core enzymatic domain. To test the specificity of the anti-vUNG antibody we performed ELISA analysis against both vUNG and murine UNG (Figure 2A). The anti-vUNG demonstrated specific binding to recombinant vUNG protein but not murine UNG. Polyclonal sera taken from the recombinant vUNG immunized mouse, displayed binding activity against both vUNG and murine UNG. In contrast, a negative control recombinant antibody, which was a mouse recombinant IgG1 antibody against the myc epitope, displayed no binding.

**Figure 1.**
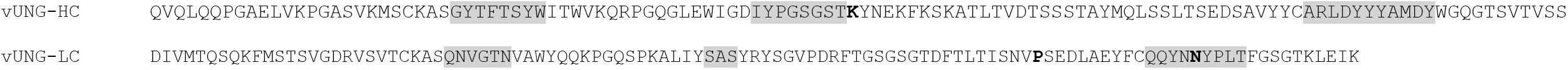
The amino acid sequence of the variable domain of the mouse immunoglobulin anti-vUNG heavy chain (HC) and light chain (LC). The sequence is encoded by the V(D)J and VJ of the chains starting from the framework 1 structure. Bold letters depict mutations from germline and the gray areas display the CDRs.

**Figure 2.**
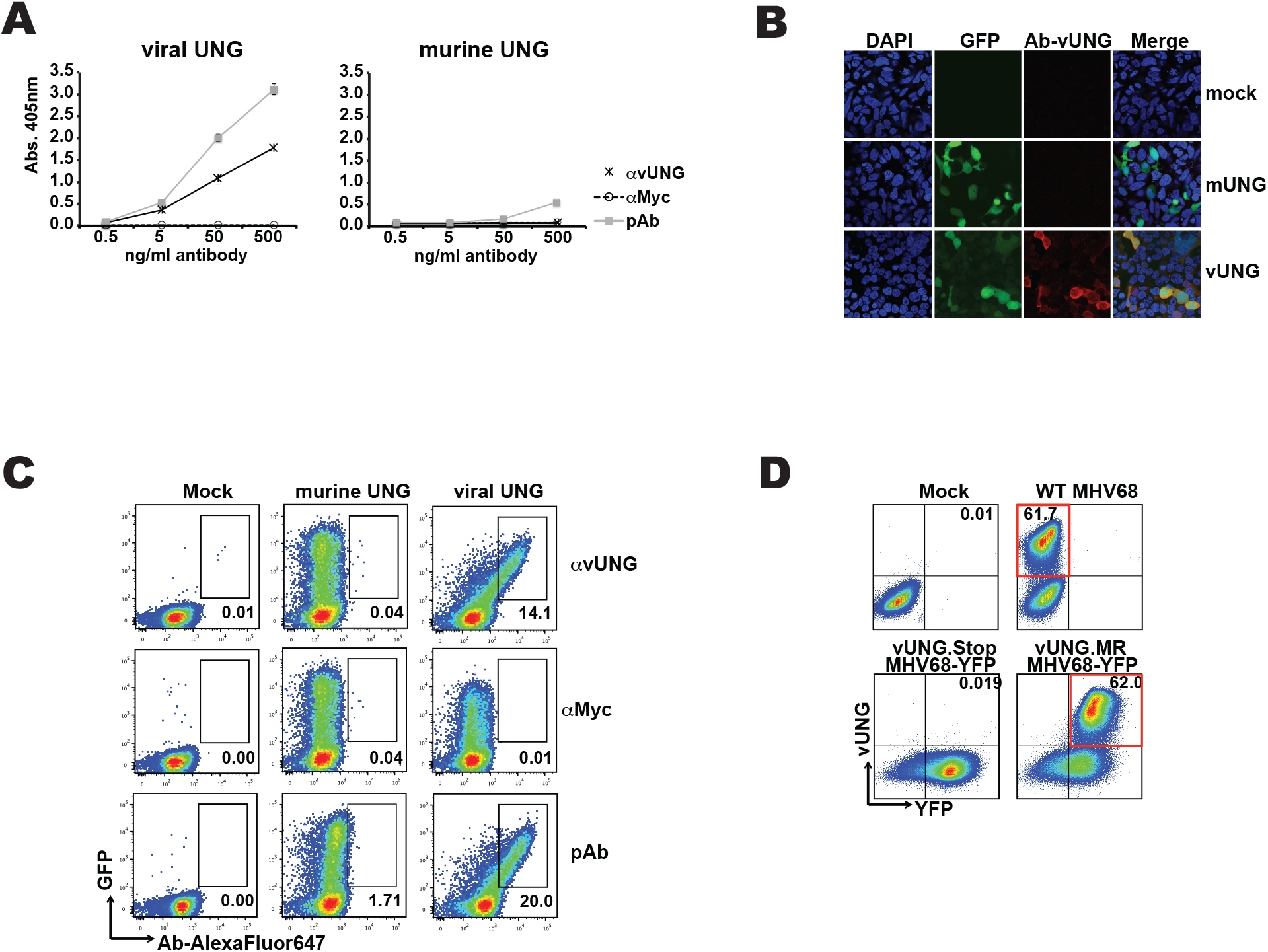
Anti-vUNG specifically detects expressed vUNG. (A) ELISA assay of anti-vUNG, anti-MYC or polyclonal sera (pAb) against purified recombinant viral UNG or murine UNG protein. Amount of antibody and OD reading is indicated on axis. (B) Microscope images of 293 cells that were mock transfected or transfected with expression plasmids encoding murine UNG (mUNG) or viral UNG (vUNG). Images for DAPI staining, GFP and antibody detection via anti-mouse (AlexaFluor647 linked) 2ndary as well as merged images are displayed. (C) Flow cytometry plots of transfected 293 cells stained with the indicated antibodies. Percentage of total cells in gates is indicated. (D) Flow cytometry plots of murine 3T12 cells mock infected, infected with MHV68 wild-type virus, MHV68-YFP with stop codon in ORF46/vUNG or MHV68-YFP marker rescue (MR) with repaired ORF46 stop codon that enables vUNG expression. Percentage of total cells in the relevant quadrant is indicated.

With the specificity to vUNG determined, we examined the suitability of the antibody to detect vUNG in expressing cells via immunostaining and microscopy or flow cytometry. We transfected HEK293 cells with a construct expressing either mUNG or vUNG and the green fluorescent protein as a transfection marker. Following fixation, permeablization and immunostaining with the anti-vUNG antibody, an anti-mouse secondary antibody coupled to Alexafluor647 was used to image the cells (Figure 2B). We detected overlap of GFP and anti-vUNG signal in cells expressing the vUNG but not in mock transfected or mUNG expressing cells (Figure 2B). For flow cytometry, the HEK293 cells were immunostained stained with the anti-vUNG antibody, polyclonal sera from immunized animals as a positive control, or a recombinant anti-myc eptitope mouse as a negative control. After flow cytometry analysis we detect overlapping signal of GFP with the anti-vUNG or polyclonal positive control sera but not the control antibody in cells expressing vUNG but not mUNG (Figure 2C). We conclude this antibody can specifically detect vUNG, but not mUNG in cells.

We next determined if the antibody could detect vUNG in cells infected with the MHV68 virus. 3T12 were harvested 24 hours post infection with wild-type MHV68 and analyzed in flow cytometry (Figure 2D). Compared to mock infection, the majority of cells displayed clear staining with anti-vUNG antibody. To show that the antibody was specifically recognizing the vUNG we used a tractable MHV68 virus that expresses Histone-H2B fused to YFP (MHV68-YFP) (Collins and Speck, 2012). We infected 3T12 cells with a mutant MHV68-YFP with a ORF-46 stop codon that disrupts vUNG expression (MHV68-YFP vUNG stop) (Minkah et al., 2015) or a marker rescue (MR) virus where the ORF-46 was repaired and vUNG expresses normally (Minkah et al., 2015) (Figure 2D). We detect no staining in the vUNG stop infected cells, but find that the majority of cells are positive for antibody staining with the repaired vUNG MR virus (Figure 2D).

We next determined if the anti-vUNG antibody could detect vUNG in immunoblots. Lysates from 293 cells either mock transfected of transfected with vUNG were run under either denaturing or native conditions. Immunoblotting was performed with either the anti-vUNG polyclonal sera or the recombinant anti-vUNG antibody. Under SDS-PAGE denaturing conditions vUNG was detected by the polyclonal sera and not the recombinant anti-vUNG antibody (Figure 3A). In contrast both the recombinant antibody and the polyclonal sera detected a band in the transfected lysates under native running conditions (Figure 3B). We conclude that the antibody can detect vUNG under non-denaturing conditions. Since the whole protein was used as the original immunogen to create the antibody, it is likely recognizing a confirmational based epitope that is disrupted under denaturing conditions.

**Figure 3.**
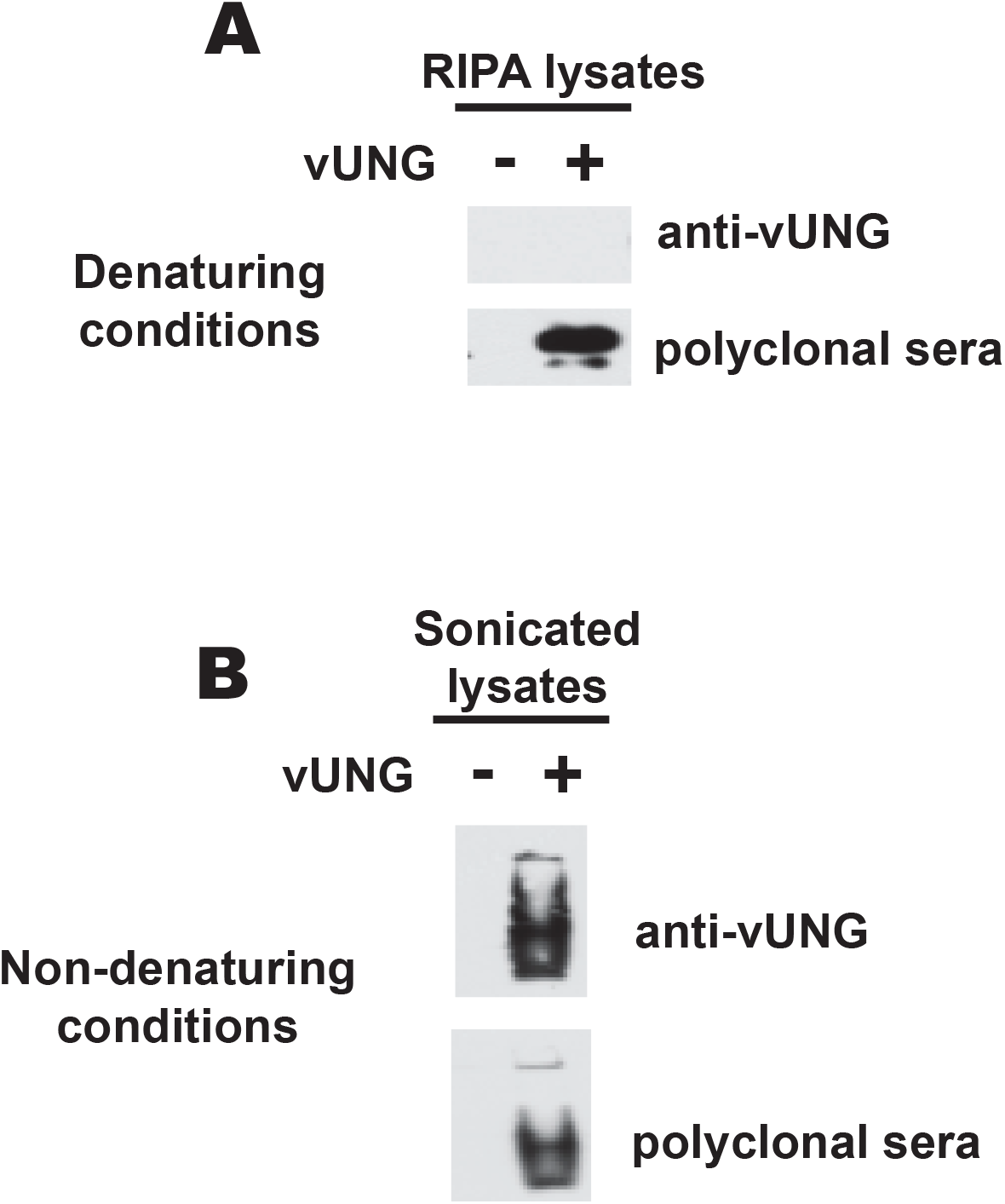
vUNG recognized by recombinant antibody under non-denaturing conditions. (A) Immunoblot of 12% NuPAGE (denaturing gel) resolving RIPA extracted cell lysates or (B) Immunoblot of 10% TBE PAGE (native gel) resolving sonicated cell lysates from 293 cells transfected with empty vector or vUNG. Blots were probed with the recombinant anti-vUNG antibody or polyclonal sera against vUNG.

In this study we characterized a new recombinant antibody against vUNG of MHV68. We clearly demonstrate that this antibody does not recognize the closely related mUNG even when overexpressed in cells lines. The anti-vUNG detects physiologic expression in virally infected murine cells and is suitable for detection in cells infected with either the wild-type or tractable MHV68-YFP virus.

## Acknowledgements

National Institutes of Health [R01AI12539 to K.M.M.; R21 Al111129 to L.T.K and K.M.M.]. Recombinant Antibody Production Core [CPRIT RP190507]. Virginia Harris Cockrell endowment fund. Texas Tobacco Settlement – Molecular Mechanisms of Tobacco Carcinogenesis. Yunxiang Mu was supported by a fellowship from the UTMDACC Center for Cancer Epigenetics.

## Conflict of interest

None declared

## Notes

### Competing Interest Statement

The authors have declared no competing interest.

